# Beauty and the brain: Investigating the neural and musical attributes of beauty during a naturalistic music listening experience

**DOI:** 10.1101/2020.10.31.363283

**Authors:** E. Brattico, A. Brusa, M.J. Dietz, T. Jacobsen, H.M. Fernandes, G. Gaggero, P. Toiviainen, P. Vuust, A.M. Proverbio

## Abstract

Evaluative beauty judgments are very common, but in spite of this commonality, are rarely studied in cognitive neuroscience. Here we investigated the neural and musical attributes of musical beauty using a naturalistic free-listening paradigm applied to behavioral and neuroimaging recordings and validated by experts’ judgments. In Study 1, 30 Western healthy adult participants rated continuously the perceived beauty of three musical pieces using a motion sensor. This allowed us to identify the passages in the three musical pieces that were inter-subjectively judged as beautiful or ugly. This informed the analysis for Study 2, where additional 36 participants were recorded with functional magnetic resonance imaging (fMRI) while they listened attentively to the same musical pieces as in Study 1. In Study 3, in order to identify the musicological features characterizing the passages that were consistently rated as beautiful or ugly in Study 1, we collected post-hoc questionnaires from 12 music-composition experts. Results from Study 2 evidenced focal regional activity in the orbitofrontal brain structure when listening to beautiful passages of music, irrespectively of the subjective reactions and individual listening biographies. In turn, the moments in the music that were consistently rated as ugly were associated with bilateral supratemporal activity. Effective connectivity analysis also discovered inhibition of auditory activation and neural communication with orbitofrontal cortex, especially in the right hemisphere, during listening to beautiful musical passages as opposed to intrinsic activation of auditory cortices and decreased coupling to orbitofrontal cortex during listening to ugly musical passages. Experts’ questionnaires indicated that the beautiful passages were more melodic, calm, sad, slow, tonal, traditional and simple than the ones negatively valenced. In sum, we identified a neural mechanism for inter-subjective beauty judgments of music in the supratemporal-orbitofrontal circuit, irrespectively of individual taste and listening biography. Furthermore, some invariance in objective musical attributes of beautiful and ugly passages was evidenced. Future studies might address the generalizability of the findings to non-Western listeners.

## 1. INTRODUCTION

That music is physical, made of sounds, signs and embodied practices, is beyond dispute. That music can be beautiful is less straightforward. On the other hand, evaluative processes of appraisal, such as beauty or liking of a song, are very common in everyday behavior. They are aesthetic concepts that are understood by “ordinary people” (Fechner 1876), that is by people that did not undertake any formal musical training. Whereas other aesthetic dimensions do exist (e.g., provocation, formal well-structuredness, originality) and might play a pivotal role for appraisal decisions in music experts, the laypersons identify the core of music aesthetics with beauty (Istók et al. 2009). Overall, beauty is a positive property attributed to an object in relation to either its physical dimensions, such as unity, harmony, balance, perfection, integrity, significant form, or to the subjective effects of the object on the perceiver, such as pleasure, satisfaction, and so on (cf. Oxford dictionary and Cambridge dictionary). The study of beauty within aesthetics has animated the philosophical debate since ancient Greece (although the term aesthetics was conceived by Baumgartner only in the 18^th^ century) and the empirical psychology research since the late 19^th^ century with Fechner’s pioneering application of psychophysics to measure aesthetic appraisal (Fechner 1876).

How musical taste, namely the set of factors guiding the positive attributions of aesthetic value to music works, develops and changes over historical epochs is the object of a whole discipline, music aesthetics (Stubley and Scruton 2002). In spite of the centrality of beauty judgments in music, empirical researchers have given much less attention to the phenomenon of musical beauty than to all other aspects of a musical object or a musical experience (whether related to listening, performing or composing). As a consequence, much less is known on what makes a song beautiful and appealing than what makes a musician a virtuoso or not. Similarly, little is known on the biological mechanisms underlying the experience of beauty in music as compared to the experience of feeling chills from a song, or being surprised by an unconventional chord (for a recent review, see Brattico, 2018).

The reasons why beauty experience is omnipresent, in all sensory-perceptual domains, are still mysterious. We humans strive in daily life to be surrounded by objects, sounds, visions, smells and even gestures to which we attribute positive qualities. Some scholars even dare to say that humans are evolutionarily adapted to seek beauty in the environment (Dutton 2009; Grammer 2018). The reasons for this have been searched for years by scholars in philosophy with the aid of speculation and theoretical formulation, then by biologists with the aid of evolutionary studies and physiology, and more recently, by psychologists and physicists with experimentation and stimulus manipulation.

With the scientific method, it is possible to explore the psychological and neural mechanisms that govern how stimuli are judged as beautiful, thus bringing new insights to a field of inquiry that has been traditionally considered mainly as the object of humanistic studies. According to Kant (Kant 2012), aesthetic judgements such as judgments of beauty are subjective judgements, but this subjectivity does not preclude the universality of these judgements or, in other words, their intersubjective agreement. Kant grounds this belief on his gnoseological theory according to which knowledge and experience are both developed through senses and “representations” that are shared among human beings. Saying it with the terms of modern neuropsychology, we share a common neurocognitive system that allows us to agree in our aesthetic judgements. This system might respond similarly to beautiful stimuli when triggered by features, functionally selected to be preferred and associated with a positive valence. Else this neurocognitive system might be activated by any feature that has been associated with a positive reaction as a result of associative conditioning or other forms of learning. Hence, when there is no agreement in aesthetic judgments, the same neurocognitive system might be sought, but the features activating it vary depending on individual factors such as expertise, cultural background, habits, etc.

The relation between the judgment of beauty and objective stimulus features as opposed to the subjective experience is a long-standing debate. In traditional musicology, this debate opposes formalists to emotivists and contextualists. The major exponent of the formalist view of musical beauty, namely the dependence of beauty on formal structural features of the music, was the German philosopher Hanslick (Hanslick and Payzant 1986). This view denies any role to emotions induced by music and rather attributes all the importance to harmony, form and other structural properties of the Western classical instrumental tradition. Here, the emotions, as well as the concepts that we usually consider as expressions of music, are instead epiphenomena and post-hoc attributions provided by humans. In this context, they would not deepen the essence of music and, on the contrary, focusing on these could be distracting, because it would bring us far away from the very essence of music.

In this study we aimed to feed this long-lasting philosophical debate by bringing new empirical evidence from cognitive neuroscience. Specifically, we used a multi-methods approach, consisting in the application of neuroimaging technology combined with a naturalistic paradigm, behavioral ratings and computational acoustic feature extraction. The focus was to empirically study the relationship between the aesthetic subjective responses to music and the intrinsic structural properties of music that determine those responses in an inter-subjectively consistent way, with the final goal of isolating the neurocognitive system that responds to musical beauty. One recent empirical effort had a similar aim but using physiological recordings. Omigie and colleagues (2019) measured the physiological bodily changes during passages of familiar tunes that were judged beautiful and tested them against subjective emotional dimensions and stimulus acoustic features. The results evidenced three distinct kinds of beauty experiences for songs, all characterized by high energy in combination with either low or high tension. Neuroimaging findings seem to support the hypothesis of an inter-subjectively consistent brain function dedicated to beauty judgments. For instance, exposure to beautiful faces (Kant 2012; Ishai 2007), paintings (Huston et al. 2015) or even architectural spaces (Huston et al. 2015; Vartanian et al. 2013) recruits the subcortical reward circuitry of the brain, which includes the ventral part of the striatum, and the ventral tegmental area. Cortically, the rewarding perception of beautiful objects recruits the ventromedial and orbitofrontal cortex, anterior cingulate, and insula.

The aim of the study was to investigate the neural correlates of aesthetic appreciation of music using a naturalistic free-listening paradigm. While some interest in research settings resembling a realistic aesthetic situation has been raised in visual cognitive neuroscience (Belfi et al. 2019), no studies have been conducted so far with music. This gap in the literature is even more astonishing when considering the temporal dynamic nature of music and of the listening experience. Most studies in the music neuroaesthetic literature addressing questions related to appraisal, enjoyment and aesthetic judgments have mostly used scarcely ecological paradigms. Typically, participants have been exposed to short and artificially modified musical excerpts or subject-selected music inside the fMRI scanner and were required to perform evaluations on discrete scales of their affective value prior, during or after the scanning session (e.g., Ishizu and Zeki 2011; Salimpoor et al. 2011; Salimpoor et al. 2011, 2013; Pereira et al. 2011; Brattico et al. 2015; for a review, see Brattico, 2019). In turn, here we wanted to record the brain neurometabolic response while participants listened attentively to three representative pieces of music of different genres, without any interruption or distraction (apart from the continuous scanner noise in the background). Continuous ratings of the perceived beauty were instead collected during a subsequent session with a different sample of participants by using a motion sensor. This procedure allowed us to identify the musical passages that would be inter-subjectively judged as more beautiful or ugly, namely consistently among all our Western participants and irrespective of their musical background knowledge.

Furthermore, we wished to determine the invariant musicological features that characterize the passages obtaining the most consistent beauty ratings by means of a post-hoc questionnaire study conducted with experts. Based on existing literature, we selected the following set of musical features: tonality, melody, complexity, novelty, and emotions expressed (joy, pathos and agitation). To our knowledge, a similar naturalistic approach in neuroimaging for studying aesthetic responses to music has been used thus far only by Trost et al. (2015), where they targeted the emotional dimensions of valence and arousal, and by Alluri et al. (2015), where liking judgments and their modulation by expertise were examined. Also, few studies have investigated the network of brain regions responding in synchrony, namely in correlation with each other, when experiencing musical emotions. The picture emerging points at two main circuits involved in emotional and aesthetic responses to music, namely the default mode network, which becomes more connected during listening to preferred music (Wilkins et al. 2014) and during listening to sad music (Alluri et al. 2015; Taruffi et al. 2017), and the auditory-limbic network linking the fronto-temporal auditory circuit to reward regions including the orbitofrontal cortex and the ventral striatum during positive appraisal of music (implicit pleasure and conscious liking of music; e.g., Liu, Abu-Jamous, et al. 2017; Liu, Brattico, et al. 2017; Alluri et al. 2015; for a review, see Reybrouck, Vuust, and Brattico 2018). Thus far, though, no neuroimaging study either examining regional activity or network connectivity has focused on musical beauty intended as a conscious aesthetic judgment operationalised with the help of continuous and discrete ratings by participants and experts.

We hypothesized to obtain significant clusters of activation during listening to musical passages concordantly rated as more beautiful across participants in regions that are related to aesthetic appraisal, such as the orbitofrontal cortex, cingulate gyrus and ventral striatum (Chatterjee and Vartanian 2016). For the passages rated as ugly we did not formulate a specific hypothesis on the areas to expect. Hence, the investigation of the brain areas activated during listening to passages that were inter-subjectively rated as ugly was mainly exploratory. However, we assumed the involvement of brain structures related to sensory processing of sounds, namely the supratemporal regions, especially for negative responses to music, and to the processing of negatively-valenced stimuli, such as the amygdala, hippocampus, parahippocampal gyrus and temporal poles (Kumar et al. 2013; Brattico et al. 2015; Blood et al. 1999; Koelsch et al. 2006; Koelsch 2018). We also predicted that the neural communication between sensory and limbic regions during listening to beautiful and ugly passages of music would be modulated by the valence of their aesthetic value, with forward and backward connections between sensory and limbic regions specifically in relation with the beauty experience.

## 2. METHOD

The study includes three separate studies conducted on three subject samples: Study 1 with behavioral measures; Study 2 with neuroimaging measures on a selection of stimuli based on results from Study 1; Study 3 with behavioral measures on the same selection of stimuli as used in Study 2. The overall aim was to investigate how intersubjectively perceived musical beauty was related to the listeners’ brain activity on the one hand, and to the structural properties of music on the other hand. The data collection for the current study was part of the broad “Tunteet” project, involving additional psychological tests, neuroimaging, neurophysiological measures and genetic mapping. The findings related to the other parts of the project are reported in separate papers (Alluri et al. 2015; Alluri et al. 2017; Burunat et al. 2016; Saari et al. 2018; Burunat et al. 2017, 2015; Bogert et al. 2016; Burunat et al. 2018; Kliuchko et al. 2018, 2016, 2015; Carlson et al. 2015). The protocol of this project was approved by the Coordinating Committee of the Helsinki and Uusimaa Hospital District, Finland, and complied with the Declaration of Helsinki. Written informed consent was obtained from all participants.

### Data and code availability statement

The data and code used in our study are available upon direct request. These terms of data and code sharing comply with the requirements of the funding body and institutional ethics approval.

### 2.1 Study 1 (behavioral)

#### 2.1.1 Participants

This study aimed at obtaining the conditions that would be used for analysis of fMRI data (further details on this study will be reported in a separate paper). Participants were 30 Western healthy adults (mainly from Finland but also other European countries), spanning a wide age range and education level, but gender balanced (age: years=29.66, *SD*=7.62, range=20-57; Education: range=high school-doctoral studies; 16 females and 14 males). Some demographic data such as education and occupation of nine subjects are missing since they chose not to report them. We chose to accept participants with a heterogeneous musical background, keeping in mind the goal of obtaining a representative and comprehensive sample of the general population. Hence, our participants had played a musical instrument from 0 to 35 years (*M*=10, *SD*= 9.33, range: 0-39). More demographic details on the participants can be found in Table 1. Upon admission to the study, participants signed an informed written consent. They were mainly recruited via university students and staff email lists and were compensated for their time spent in the lab and traveling with culture and sport vouchers.

**Table 1.** Demographic information of participants for Study 1. Data were collected through questionnaires. Hand. = handedness (R=right, L=left); Years = number of years of musical practice; F = female; M = male.

| **STUDY 1 (Behavioral)** |  |  |  |  |  |  |
| --- | --- | --- | --- | --- | --- | --- |
| **Subject** | **Age** | **Hand.** | **Sex** | **Education** | **Occupation** | **Years** |
| 1 | 29 | R | F | High school | Bachelor student/musician | 23 |
| 2 | 27 | R | F | Bachelor degree | Research assistant/musician | 20 |
| 3 | 34 | R | F | Master degree | Doctoral student/musician | 28 |
| 4 | 22 | R | F | Bachelor degree | Master student | 10 |
| 5 | 27 | R | M | Master degree | Doctoral student | 10 |
| 6 | 23 | R | F | Missing | Missing | 7 |
| 7 | 21 | R | M | High school | Student | 14 |
| 8 | 23 | R | M | Bachelor degree | Master student | 8 |
| 9 | 27 | R | F | Master degree | Trainee | 18 |
| 10 | 36 | R | F | Master degree | Economist | 1 |
| 11 | 20 | L | F | High school | Student | 2 |
| 12 | 29 | R | M | Bachelor degree | Student | 11 |
| 13 | 27 | R | F | Bachelor degree | Master student | 0 |
| 14 | 30 | R | F | High school | System support | 17 |
| 15 | 33 | R | M | Bachelor degree | Student | 0 |
| 16 | 31 | R | F | Master degree | Student | 0 |
| 17 | 28 | R | M | Master degree | Engineer | 0 |
| 18 | 57 | R | F | Master degree | Teacher | 11 |
| 19 | 32 | R | M | Master degree | Doctoral student | 8 |
| 20 | 33 | R | F | Missing | School teacher | 2 |
| 21 | 30 | R | M | Missing | Entrepreneur | 6 |
| 22 | 39 | R | F | Master degree | Doctoral student | 0 |
| 23 | 29 | R | M | Bachelor degree | Master student | 0 |
| 24 | 38 | R | M | Bachelor degree | Teacher | 4 |
| 25 | 43 | R | M | Music university | Musician/technician | 35 |
| 26 | 25 | R | F | Master degree | Doctoral student | 17 |
| 27 | 27 | R | M | Master degree | Doctoral student | 21 |
| 28 | 26 | R | M | Master degree | Doctoral student | 14 |
| 29 | 24 | R | M | Master degree | Doctoral student | 13 |
| 30 | 20 | R | F | High school | Student | 0 |

#### 2.1.2 Materials and procedures

Three musical pieces were employed in the study (Figure 1): “Adios Nonino” by the Argentinian composer Astor Piazzolla (1959) and “Rite of Spring” (comprising the first three episodes from Part I: Introduction, The Augurs of Spring: Dances of the Young Girls and Ritual of Abduction) by the Russian born composer Igor Stravinsky (1947), and “Stream of Consciousness” by Dream Theater (2003). In the following, the pieces will be referred to simply as Piazzolla, Stravinsky, Dreamtheater. These pieces were an Argentinian New Tango, an iconic 20th century classical work and a progressive rock piece, respectively. These pieces of music were chosen based on the following criteria: to have an appropriate duration for the research setting, to belong to different genres in order allow generalization of the obtained findings, to contain a high amount of acoustic variation and that the amount be comparable between the pieces, to have a comparable musical structure (starting with a session of solo instrument and then introducing the larger ensemble after few minutes), and lastly, to not contain lyrics in order to avoid the confounding effects of semantics (Alluri et al., 2015; Brattico et al., 2011).

**Figure 1.**
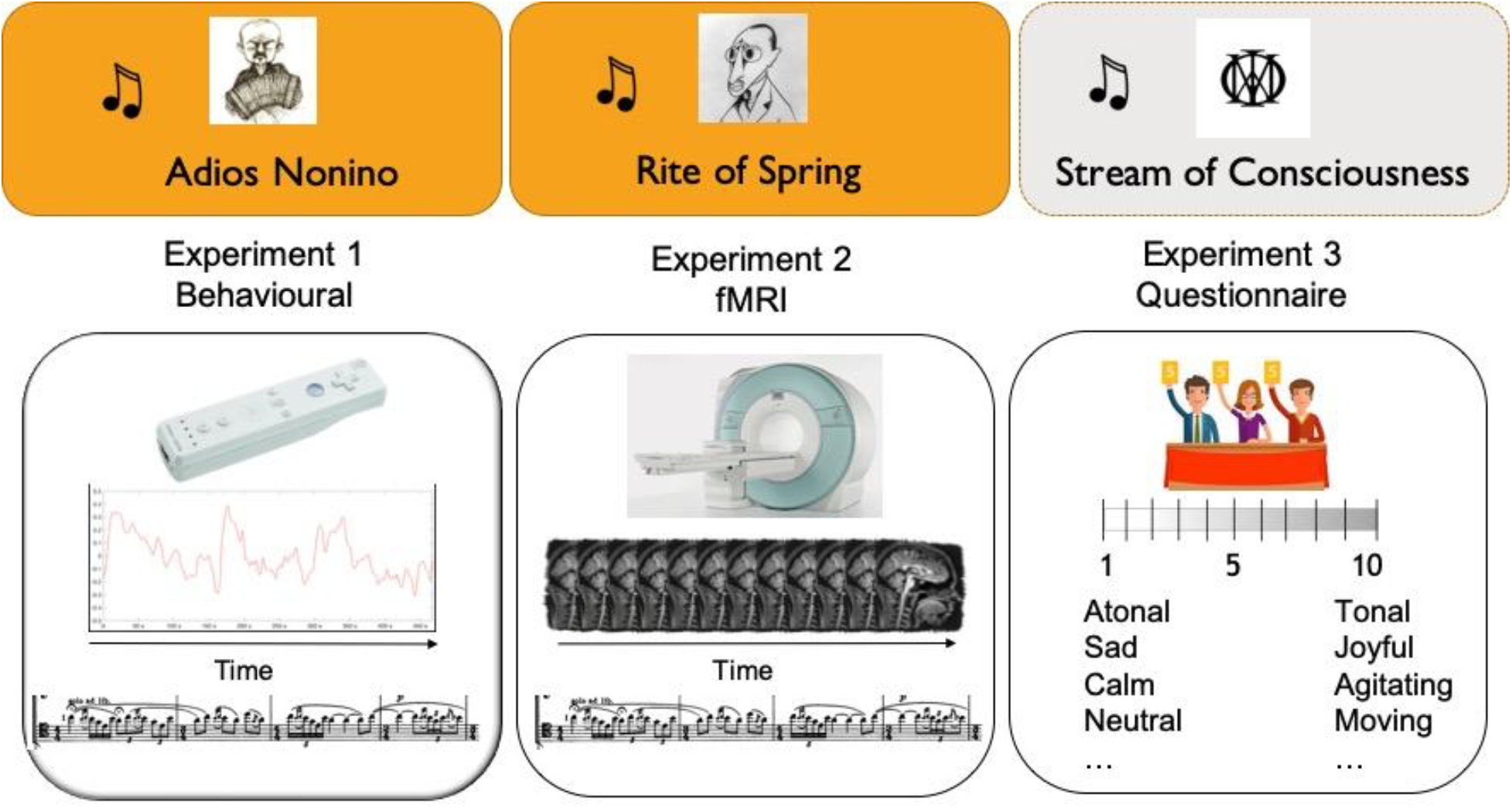
Schematic illustration of the protocol designed for this study, including 3 studies with 3 separate groups of participants, amounting to a total of 78 participants. In all studies the same three musical stimuli were used: *Adios Nonino* by A. Piazzolla, *Rite of Spring* (the first 3 dances) by I. Stravinski, *Stream of Consciousness* by Dreamtheater.

Data were collected at four different locations in Finland, all comparable in terms of sound background, equipment and experimenters coordinating data collection (Cognitive Brain Research Unit and Biomag laboratory, University of Helsinki; Department of Music, University of Jyväskylä; MIB, Aarhus University).

In this study, participants performed continuous evaluations of the perceived beauty and ugliness of the three-music tracks, presented in counterbalanced order, by using a Nintendo Wii motion sensor. The sensor allowed them to indicate over time the level of beauty subjectively experienced by moving it vertically while listening to the musical pieces. Participants received the following instructions: “raise the Wii when you find the music beautiful and lower it when you find it ugly”. Afterwards, participants also gave some discrete ratings including aesthetic preference (liking) and familiarity on a Likert scale from 1 to 5. Notably, participants with different levels of musical expertise did not differ in their liking or familiarity ratings for the three pieces, as verified with independent samples t-tests (p>.2). The collected Wii data were recorded at a frequency of 2 Hz using WiiData Capture (Version 2.2, 2012, Petri Toiviainen, Brigitta Burger, University of Jyväskylä, Finland). Inside the program, the OSCulator software was used to receive data from the Nintendo Wii remote control via Bluetooth and to prepare them to be used in WiiDataCapture.

#### 2.1.3 Data analysis

The continuous behavioral ratings were preprocessed by means of MoCap toolbox (Toiviainen, Burger, 2015, MCT Manual v1.5) and custom-made scripts in MATLAB environment (version R2106b, MathWorks, Natick, MA, USA). First MoCap was applied to read the data files saved with the WiiDataCapture application, parse them and interpolate them to 10ms intervals. This process returned a data structure containing the recorded locations of the remote in three columns, corresponding to the two horizontal dimensions and the vertical dimension. For this study, only the y-axes coordinates have been taken into consideration. The signal of each participant was normalized by the individual maximum value and centered by the mean. To visualize the data of each participant, the string of y-coordinates was plotted on a line graph where the y-axis indicated the recorded position of the motion sensor and the x-axis represented the time course of the music (see Figure 2). Subsequently, the continuous rating data were analyzed using custom-made scripts using inter-subject correlation (ISC). The goal was to identify the fragments of music that were most consistently evaluated as beautiful and ugly across participants. Hence, for each music, we firstly considered the individual signals corresponding to the aesthetic evaluation given by each participant to each music track (Figure 1). Therefore, we transformed each individual beauty rating into a binary signal. In this way, we treated the signal as positive when it was above the zero and negative when it was below it. For each time point, we summed the binary signals of the overall subjects in a single vector. We considered an agreement of 100 % in the evaluation of each time point as positive or negative when the sum of the whole subjects was, respectively, above or below zero in that specific time point. Since just few time points had a 100% agreement, we decided to lower the threshold to 70%. Subsequently, the time intervals that had at least a 70% agreement between participants and that had an average beauty signal value higher than 0.15 were selected as “Beautiful” passages, while those with a 70% agreement and an average value lower than −0.15 where selected as “Ugly” passages. The selection of all passages was then revised by two expert listeners (one of them coauthor of the paper: GG) through a careful music listening, in order to verify the consistency between the music and the ratings (and in order to assess the initial delay of participants’ movements for an interval they wanted to rate as beautiful or ugly). The final selection of beautiful and ugly passages from Piazzolla and Stravinsky musical pieces is visible in Table 2:The ratings for Piazzolla consisted of 48750 values, corresponding to the 487.968 seconds of the music, Dreamtheater of 47100 values, corresponding to 471.250 seconds of the music and Stravinski of 46750 values, corresponding to 472.034 seconds of the music (see Table 4).

**Table 2.** Selected musical passages derived from the analysis of behavioral ratings of beauty in Study 1. For each musical piece two sets of time intervals were extracted; one representing the intervals rated as beautiful (beautiful passages), and one representing the ones rated as ugly (ugly passages). Overall, the passages for the Piazzolla piece comprised 98 seconds in the beauty condition, and 95 seconds in the ugly condition; for the Stravinsky piece they lasted in total 74 and 82 seconds, respectively.

| Piazzolla |  | Stravinsky |  |
| --- | --- | --- | --- |
| Beautiful passages | Ugly passages | Beautiful passages | Ugly passages |
| 01:30.000-01:50.000 | 00:36.000-01:08.000 | 00:09.000-00:43.000 | 01:55.000-02:10.000 |
| 03:50.000-04:23.000 | 01:57.000-02:12.000 | 02:50.000-03:10.000 | 02:34.000-02:46.000 |
| 04:37.000-04:52.000 | 02:55.000-03:08.000 | 05:02.000-05:12.000 | 03:35.000-03:45.000 |
| 06:22.000-06:40.000 | 05:40.000-06:15.00 | 05:22.000-05:42.000 | 03:55.000-04:05.000 |
| 07:08.000-07:20.000 |  |  | 04:40.000-04:55.000 |
|  |  |  | 06:30.000-06:40.000 |
|  |  |  | 07:16.000-07:26.000 |

**Figure 2.**
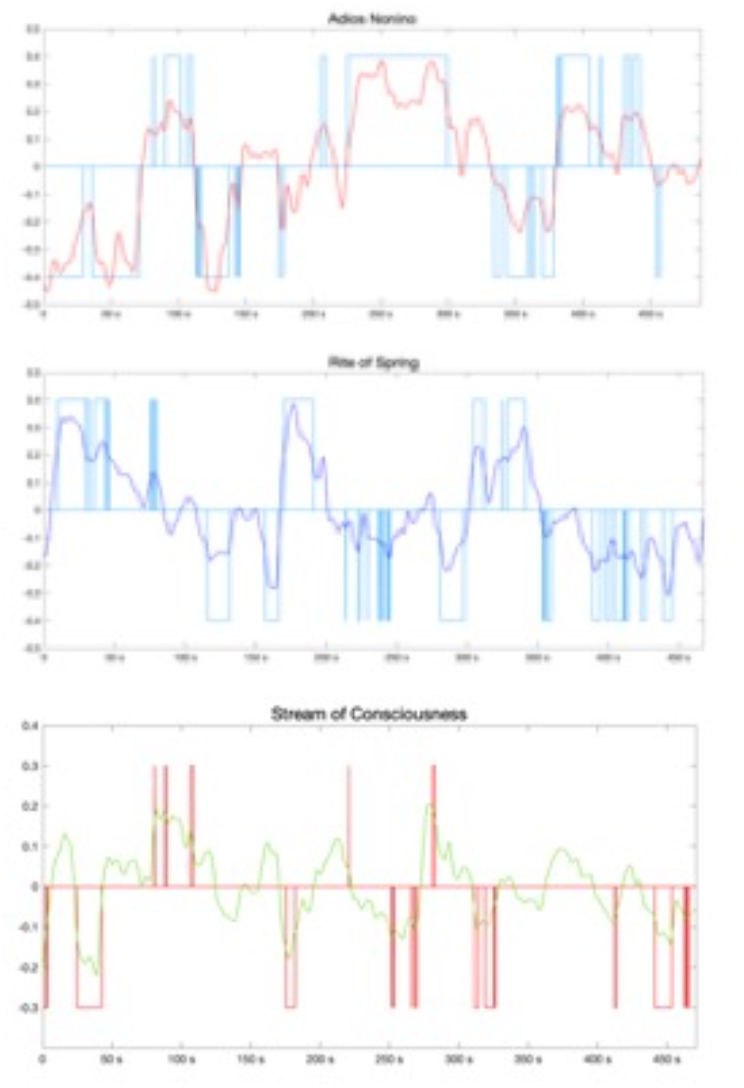
Average continuous ratings of beauty over the music time course from 30 participants during listening to the three musical stimuli used in Study 1: *Adios Nonino* by Piazzolla (top), *Rite of Spring* (the first 3 dances) by Stravinski (middle), *Stream of Consciousness* by Dreamtheater (bottom). The rectangles indicate the musical passages that were most consistently rated positively (beautiful) or negatively across participants of Study 1 (70% for Piazzolla and Stravinski and 65% for Dreamtheater).

### 2.2 Study 2 (fMRI)

#### 2.2.1 Participants

Thirty-six (n=36) healthy participants took part in an fMRI session which took place at the Advanced Magnetic Imaging (AMI) Center of Aalto University School of Science, Espoo, Finland. Participant’s pool covered a wide spectrum of ages and education and was gender balanced (age: years=28.75, *SD*=9.21, range=18-52; Education: range=secondary school-doctoral degree; 15 Females and 17 Males). Few demographical data such as education and occupation of nine subjects are missing since they chose not to report them. The participants had no history of neurological or psychiatric diseases. They were screened for inclusion criteria before admission to the Study (no ferromagnetic material in their body, no tattoo or recent permanent coloring, no pregnancy or breastfeeding, no chronic pharmacological medication, no claustrophobia) and upon admission they signed an informed written consent. Subjects were mostly recruited via university students and staff email lists, they were compensated for their time spent in the lab and traveling in the form of culture vouchers that could be used for cultural and sport activities (e.g., for going to a swimming pool, or buying a museum ticket).

As evidenced in Table 3, participants had a heterogeneous experience and expertise in music as shown by the amount of years they have been playing a music instrument (*M*=11.65, *SD*= 11.28, range: 0-39). All participants considered music moderately important in their lives (*M*=5.50, *SD*=1.46 on a Likert scale from 1 to 7). Detailed information can be found in Table 3. The sample of participants was comparable for demographic variables with the sample from Study 1: between-group paired t-tests for age: t(64)=.43, p=.66; years of playing an instrument: t(64)=-.65, p=.52).

**Table 3.** Demographic information on the participant of Study 2. Data were collected through questionnaires. Hand.= handedness (R=right, L=left); Years = number of years of musical practice; F = female; M = male.

| STUDY 2 (fMRI) |  |  |  |  |  |  |
| --- | --- | --- | --- | --- | --- | --- |
| Subject | Age | Hand. | Sex | Education | Occupation | Years |
| 1 | 27 | R | F | Bachelor's degree | Student | 0 |
| 2 | 20 | R | F | Missing | Missing | 15 |
| 3 | 18 | R | F | Missing | Missing | 13 |
| 4 | 20 | R | M | Secondary school | Student | 16 |
| 5 | 33 | R | M | Bachelor's degree | Musician | 21 |
| 6 | 21 | R | M | Secondary school | Student, freelance musician | 17 |
| 7 | 39 | R | F | Missing | Missing | 30 |
| 8 | 19 | R | M | Secondary school | Student | 0 |
| 9 | 25 | R | F | Missing | Missing | 20 |
| 10 | 34 | R | M | PhD | Researcher | 11 |
| 11 | 31 | R | M | Master's degree | Musician | 25 |
| 12 | 23 | R | F | Secondary school | Student | 0 |
| 13 | 44 | R | M | Music graduate | Musician | 39 |
| 14 | 40 | R | M | Master's degree | Musician/homemaker | 30 |
| 15 | 28 | R | F | Bachelor's degree | Master student | 0 |
| 16 | 21 | R | F | Secondary school | Student | 0 |
| 17 | 27 | R | F | Bachelor's degree | Research assistant | 20 |
| 18 | 34 | R | F | Master's degree | Doctoral student | 28 |
| 19 | 24 | R | M | Missing | Missing | 0 |
| 20 | 26 | R | M | Missing | Missing | 2 |
| 21 | 21 | R | M | Secondary school | Student | 2 |
| 22 | 35 | R | F | Bachelor's degree | Piano teacher | 29 |
| 23 | 25 | R | M | Master's degree | Research assistant | 11 |
| 24 | 33 | R | F | Master's degree | Student | 4 |
| 25 | 24 | R | M | Bachelor's degree | Student/research assistant | 19 |
| 26 | 22 | R | F | Bachelor's degree | Master student | 10 |
| 27 | 23 | R | F | Missing | Missing | 7 |
| 28 | 28 | R | F | Master's degree | Doctoral student | 5 |
| 29 | 29 | R | F | Music bachelor | Composer | 24 |
| 30 | 52 | R | M | Master's degree | Researcher | 4 |
| 31 | 50 | R | F | Master's degree | Foreign language teacher | 0 |
| 32 | 20 | R | F | Missing | Missing | 0 |
| 33 | 29 | L | M | Missing | Missing | 2 |
| 34 | 19 | R | M | Secondary school | Student | 0 |
| 35 | 49 | R | F | Master's degree | Assistant | 2 |
| 36 | 21 | R | M | Secondary school | Student | 14 |

#### 2.2.2 Materials and procedures

The fMRI study adopted the *naturalistic free-listening* paradigm (Alluri et al. 2012) where participants were asked to listen attentively to entire musical pieces without any interruption. The musical stimuli were the same as used in Study 1 and similarly, they were presented in counterbalanced order. Prior to the start of the MR session, subjects filled screening questionnaires and were given instructions on the study session. Subsequently, they entered the MR scanner, were given high-quality MR-compatible insert earphones (custom-made at AMI Center) and then the stimulus loudness was adjusted to a comfortable but audible level inside the scanner room (around 75 dB) by presenting to the subjects an acoustically comparable stimulus to the ones used in the study. Then, participants were asked to lay still, with their eyes open inside the scanner, and listen attentively to the music.

During fMRI recordings, listening was not interrupted by any intermittent behavioral or motoric task. However, listeners were aware that after each piece they would be asked some questions on the music, namely Likert ratings on a scale from 1 to 5 including familiarity and liking. This procedure was aimed at maintaining their attention on listening and also on acquiring discrete ratings for each musical piece in the course of the fMRI session (tested with paired-samples Wilcoxon sign-rank tests). Post-session debriefing confirmed their engagement in the listening experience.

The order of presentation of the musical pieces was counterbalanced across participants and each piece was followed by 1-2 minutes break during which the experimenter asked questions related to the stimulus ratings via the interphone and marked the answers on a paper. The collection of functional EPI images lasted in total around 30 minutes including breaks. After EPI recordings, participants remained inside the MR-scanner for structural (MPRAGE) images collection, lasting around 15 minutes in total.

#### 2.2.3 fMRI data acquisition and analysis

Brain scanning was performed using a 3T MAGNETOM Skyra whole-body scanner (Siemens Health-care, Erlangen, Germany) and a standard 20-channel head-neck coil, at the Advanced Magnetic Imaging Centre (Aalto University, Espoo, Finland). Using a single-shot gradient echo planar imaging (EPI) sequence thirty-three oblique slices (field of view = 192×192 mm; 64×64 matrix; slice thickness = 4 mm, interslice skip = 0 mm; echo time = 32 ms; flip angle = 75°) were acquired every 2 seconds, providing whole-brain coverage. T1-weighted structural images (176 slices; field of view = 256×256 mm; matrix = 256×256; slice thickness = 1 mm; interslice skip = 0 mm; pulse sequence = MPRAGE) were also collected. Functional MRI scans were preprocessed on a Matlab platform using SPM8 (Statistical Parametric Mapping; https://www.fil.ion.ucl.ac.uk/spm/software/spm8/), and VBM5 for SPM (Voxel Based Morphometry; Wellcome Centre for Human Neuroimaging, London, UK; http://dbm.neuro.uni-jena.de/vbm/), in order to remove the skull before proceeding to the normalization step. In more detail, for each participant, low-resolution images were realigned on six dimensions using rigid body transformations (translation and rotation corrections did not exceed 2 mm and 2° respectively), segmented into grey matter, white matter, and cerebrospinal fluid, and registered to the corresponding segmented high-resolution T1-weighted structural images. These were in turn normalized to the MNI (Montreal Neurological Institute) segmented standard a priori tissue templates using a 12-parameter affine transformation. Functional images were then smoothed to best accommodate anatomical and functional variations across participants as well as to enhance the signal-to-noise by means of an 8-mm full-width-at-half-maximum Gaussian spatial filter. Movement-related variance components in fMRI time series resulting from residual head-motion artifacts, assessed by the six parameters of the rigid body transformation in the realignment stage, were regressed out from each voxel time series in subsequent analyses.

Subsequently, the fMRI data were analysed using General Linear Model (GLM) with SPM12 (Statistical Parametric Mapping; http://www.fil.ion.ucl.ac.uk/spm/software/spm12/). At first-level functional images the music fragments obtained from Study 1 were modelled as single regressor including two conditions: “Beautiful” and “Ugly”, by entering the respective list of onsets with different durations. The rest of the music was treated as baseline. Each block was at least 10 seconds long and the intervals between selected trials were heterogeneous, different each time.

In an additional GLM analysis, we also included as regressors of no interest the same six acoustic components as obtained with the MIRToolbox (Lartillot, Toiviainen, and Eerola 2008) in (Vinoo Alluri et al. 2012) of Fullness, Brightness, Activity, Timbral Complexity, Key Clarity and Pulse Clarity for each music piece in order to eliminate the variance generated by the sensory processing of them.

In both GLM analyses, stimulus functions were convolved with a canonical double gamma Hemodynamic Response Function (HRF). Beta parameters were estimated and the contrast images beautiful>baseline and ugly>baseline were generated. These contrasts were then taken to second-level (between-subject) analysis to produce statistical parametric maps at the group level. For both GLM analyses (with or without acoustic components regressed out), at this second-level we pruned from the brain signal the variance accounted for by the familiarity to the pieces, inserting the discrete values of familiarity for each participant and each musical piece as covariate. For both the GLM analyses, we used a factorial design which consisted of two factors: (a) Musical Piece (Piazzolla and Stravinsky) and (b) Aesthetic Value (beautiful and ugly).

#### 2.2.4 Analysis of effective connectivity

We further wished to test understand whether the aesthetic judgments would modulate the directionality in connections between auditory cortex and emotion-related brain structures. For this analysis, we started from the GLM results and isolated the brain regions activated during listening to beautiful versus ugly music. Then, we designed effective connectivity models by means of dynamic causal modelling (DCM). The methodological details of this modelling are reported below.

##### Volumes of interest (VOIs)

We first extracted BOLD time-series from three volumes-of-interest (VOIs) in each participant using the first principal component of voxels within a sphere of 8 mm radius centred on each participant’s local maximum. The subject-specific local maximum was identified within a sphere of 20 mm radius centred on the peak of the group effect identified the random-effects GLM analysis. Only voxels exceeding a threshold of p < 0.05, uncorrected were included in each VOI. One subject was excluded from the DCM analysis due to the lack of signal in the left STG.

##### Dynamic causal modelling of effective connectivity

We used a two-state DCM for fMRI (DCM12, revision 7487) to estimate the effective connectivity within and between brain areas, given observed hemodynamic measurements. Two-state DCM models both extrinsic connections between regions as excitatory forward and backward connections and intrinsic connectivity within each region in terms of one inhibitory and one excitatory population of neurons. These models allowed us to portrait the intrinsic connectivity within each cortical area as an increase or decrease in cortical inhibition (Marreiros, Kiebel, and Friston 2008). The network deriving empirically from the group-level GLM results comprised bilateral superior temporal gyrus (STG) and medial orbito-frontal cortex (OFC) (for details on GLM findings see Results section). Hemodynamic responses to all auditory stimuli were modelled as a driving input to left and right STG. Using parametric modulation of the regressor encoding all auditory stimulus blocks, responses to ‘beautiful’ stimuli compared to ‘ugly’ stimuli were modelled as a modulatory input to the network under four alternative hypotheses, all illustrated in Figure 5A. The first hypothesis was formulated as a full model where both extrinsic (excitatory) connections between bilateral STG and OFC and intrinsic (inhibitory) connections within each region encode the difference between the value of ‘beauty’ and the value of ‘ugliness’. The second hypothesis was formulated as a reduced model where only extrinsic (excitatory) connections between bilateral STG and OFC encode the differences between conditions. The third hypothesis was formulated as another reduced model where only the forward (excitatory) connections from bilateral STG to OFC encode the differences between conditions. Finally, the fourth hypothesis tests the belief that no connections encode any differences between conditions. We then used Bayesian model reduction (BMR) to estimate the posterior probability of the connection strengths and the Bayesian model evidence of each alternative model within each subject (Friston et al. 2016).

**Figure 5.**
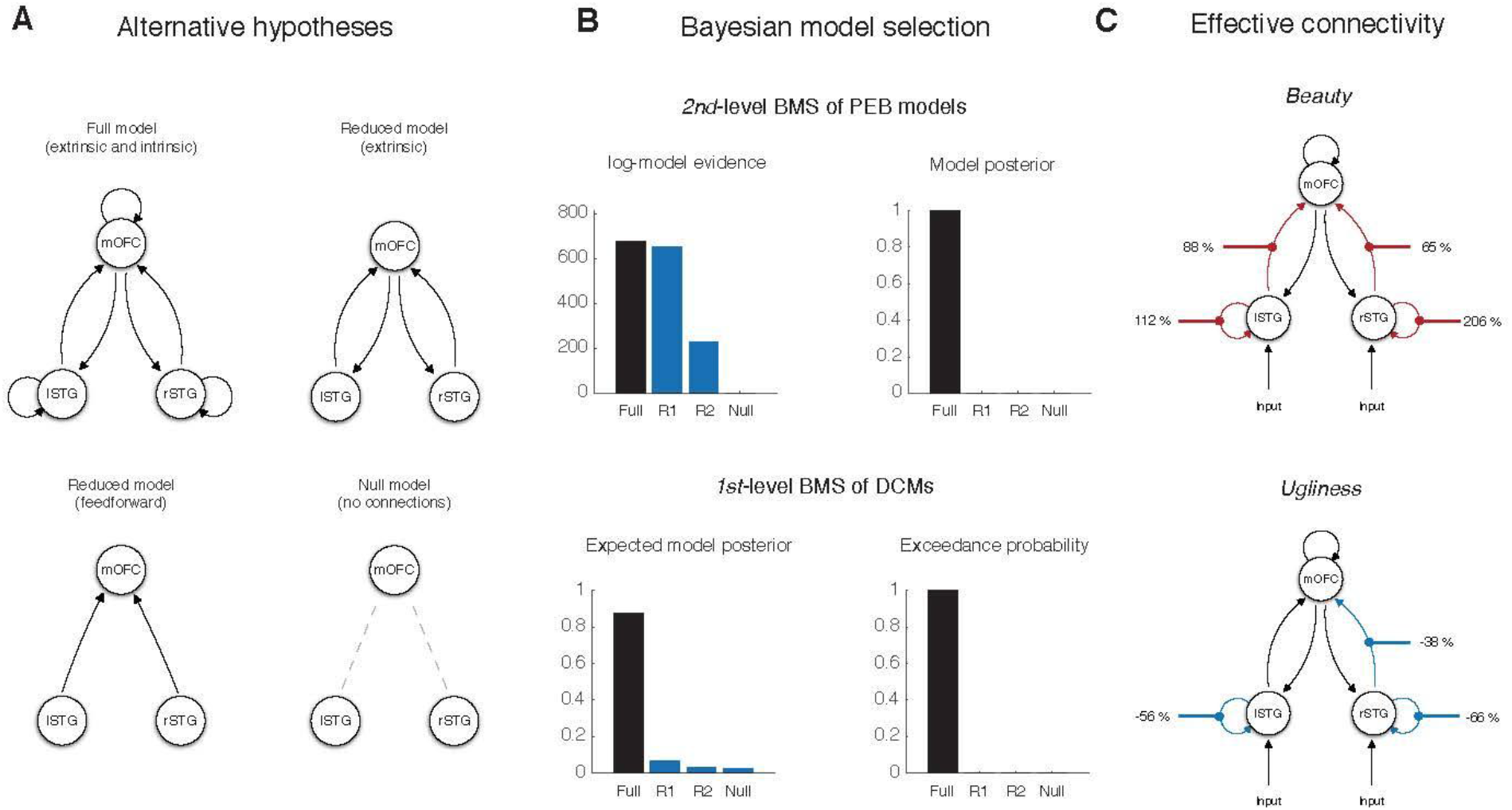
Dynamic causal modelling of effective connectivity **a)** Alternative hypotheses about effective connectivity during listening to beautiful or ugly musical passages. **b)** Bayesian model selection of DCMs at the first level showing posterior model probabilities and protected exceedance probabilities and Bayesian model selection of PEB models at the second level showing log-evidences and posterior model probabilities. **c)** Graphical structure of the DCM with the highest model evidence showing a bilateral increase in forward connectivity and a concomitant increase in STG inhibition during listening to beautiful passages of music and a decrease in right-lateralized forward connectivity accompanied by a decrease in STG inhibition.

##### Parametric Empirical Bayesian analysis of group effects

We used parametric empirical Bayes (PEB) to identify increases or decreases in extrinsic (excitatory) connections between bilateral STG and OFC and intrinsic (inhibitory) connections within each region at the group level. PEB is a hierarchical Bayesian model in which empirical priors on the connection strengths at the single-subject level are estimated empirically using a Bayesian general linear model at the group level (Friston et al. 2016).

### 2.3 Study 3 (questionnaire)

#### 2.3.1 Participants

Twelve (n=12) music composition experts were engaged for this part of the research (demographic details can be found from Table 4), in order to determine whether the beautiful and ugly passages identified from Study 1 and used in Study 2 would be marked by a specific set of invariant musicological features, as evidenced by ratings obtained from music-composition experts based in Milan. The group comprised seven renowned music composers and conservatory professors of composition, two orchestra directors and three conservatory students of composition. The experts included three women and nine men aged between 20 and 74 years old (*M*=47.25, *SD*=17.44). They were recruited on voluntary base through local conservatory and mailing lists and were compensated for their time spent with a twenty euros Amazon voucher. More information on the participant sample can be found in Table 4.

**Table 4.** Demographic information of participants for Study 3. Data were collected through paper-and-pencil questionnaires. F = female; M = male.

| STUDY 3 (Questionnaire) |  |  |  |  |
| --- | --- | --- | --- | --- |
| Subject | Age | Sex | Education | Occupation |
| 1 | 20 | M | Undergraduate at Conservatory | Conservatory student |
| 2 | 21 | M | Undergraduate at Conservatory | Conservatory student |
| 3 | 55 | M | Conservatory | Music composer, Professor of composition |
| 4 | 50 | M | Conservatory | Music composer, Professor of composition |
| 5 | 57 | M | Conservatory | Music Composer, Professor of composition |
| 6 | 56 | M | Conservatory | Music Composer, Professor of composition |
| 7 | 20 | M | Undergraduate at Conservatory | Conservatory student |
| 8 | 47 | F | Conservatory | Orchestra director |
| 9 | 74 | F | Conservatory | Professor of composition |
| 10 | 56 | M | Conservatory | Music composer, Professor of composition |
| 11 | 57 | M | Conservatory | Music composer, Professor of Composition |
| 12 | 54 | F | Conservatory | Orchestra director |

#### 2.3.2 Stimuli

Stimuli were the “Beautiful” and “Ugly” musical passages as derived from Study 1 and selected as regressors for the fMRI-data analysis of Study 2.

#### 2.3.3 Procedures

Music composition experts were asked to evaluate on a list of aesthetic and musical features each fragment of music used as conditions in the fMRI analyses. Participants completed an on-line or equivalent paper-and-pencil questionnaire prepared ad hoc for this study where they had to indicate the tonality of each excerpt and to evaluate each one of them on a dimensional scale from 1 to 10 on the following dimensions: a) atonal-tonal, b) neutral-moving, c) calm-agitating, d) sad-joyful, e) slow-fast, f) rhythmic-melodic, g) simple-complex (rhythmically), h) simple-complex (harmonically), i) simple-complex (performance), j) boring-interesting, k) trivial-original, l) prosaic-sublime, m) traditional-innovative. They had to indicate whether the excerpt was better represented by either one of the poles of each dimension by assigning a value closer to that end. For instance, considering the dimension atonal-tonal, when the excerpt was evaluated as atonal they had to assign a value closer to 1, on the contrary when it was atonal the value was closer to 10.

To evaluate the excerpts, they received a Dropbox link to the same audio files used in the other parts of the study, corresponding musical scores of the pieces, and a table indicating the beginning and end of each musical passage to be evaluated. Participants were kept in the dark concerning the value (beautiful/ugly) assigned to the selected intervals of music. They were instructed to listen to the fragments of music one by one, examine the music score corresponding to them and evaluate them. The questionnaire took around 1,5 hours to complete.

#### 2.3.4 Data analysis

The data collected were analyzed using SPSS software (IBM SPSS Statistics for Macintosh, Version 24.0). Separate analysis of Variance (ANOVA) tests were performed with a factorial design to investigate the extent to which ‘beautiful’ musical excerpts differed from the ‘ugly’ passages on each aesthetic dimension and each musical piece. The scores assigned to one aesthetic dimension in the questionnaire were used as *within group* dependent variable, while the Aesthetic Value (two levels: beautiful and ugly) and the Musical Piece (two levels: Piazzolla and Stravinsky) were used as *between group* factors. The analysis was repeated for each aesthetic dimension of the questionnaire.

## 4. RESULTS

### 4.1 Study 1

#### 4.1.1 Discrete behavioral ratings

##### Aesthetic preferences

The repeated measures ANOVA carried on the discrete liking ratings on a scale from 1 to 5, where 1 indicated “not liking at all” and 5 “liking very much”, did not show any significant main effect of the musical piece (F=1). Piazzolla was liked on average M=3.6, SD=1.3, Stravinsky M=3.23, SD=1.31, and Dreamtheater liking scores of M=3.2, SD=1.49.

##### Familiarity

The analyses showed that, on a Likert scale from 1 to 5, where 1 indicated “not familiar at all” and 5 indicated “very familiar”, familiarity ratings did not differ according to the musical pieces (F<1). Piazzolla received an average familiarity score on M=3.03 (SD=1.33), Stravinski of M=2.87 (SD =1.43), while Dreamtheater of M=2.57 (SD =1.361), indicating that listeners on average were only mildly familiar with those pieces.

#### 4.1.2 Intersubject-correlations of continuous ratings

The average signal of the continuous ratings collected with Wii motion sensor is shown in Figure 2. The passages that were most consistently (over 70% agreement) rated as beautiful or ugly by participants were identified as in Table 2. Those passages were then used for constructing the regressor for Study 2. The music piece by Dreamtheater was excluded at this point from further analyses since agreement across raters’ aesthetic evaluations did not reach our threshold set at 70% (following to standards used for using other reliability measures such as Cronbach’s alpha). A separate study containing all the results obtained for this behavioral study will be reported in a separate paper.

### 4.2 Study 2

#### 4.2.1 Discrete behavioral ratings

##### Aesthetic preferences

The repeated measures ANOVA carried on the liking ratings in a scale from 1 to 5, where 1 indicated “not liking at all” and 5 “liking very much”, showed a significant main effect of the musical piece (F(2,70)=9.34, p<0.001). Tukey post-hoc test indicated that while ratings for Piazzolla and Stravinsky did not differ between participants (Piazzolla: M=3.83, SD=0.97; Stravinski: M=3.58, SD=1.25) Dreamtheater received lower liking scores (M=2.75, SD=1.18) than both Piazzolla and Stravinsky (p<0.03). However, when comparing the musical preferences of participants from Studies 1 and 2, we did not yield any main effect of Study (F<1) nor any interaction with Musical Piece (F(1,128)=2.27, p=.11), suggesting a general lower enjoyment of the Dreamtheater piece as compared with the other two. Moreover, when pooling participants from both Studies 1 and 2, correlation analyses showed that the amount of musical practice correlated with the degree of appreciation for Stravinsky’s musical piece (r=0.46, p<.01) but not for Piazzolla’s (r=0.24, p=.16), nor for Dreamtheater’s (r=-.28, p=.09).

##### Familiarity

The analyses showed that familiarity ratings did not differ according to the musical pieces (F(2,70)=2.65, p=0.08). Piazzolla received an average familiarity score on M=2.75 (SD=1.36), Stravinsky of M=2.75 (SD =1.65), while Dreamtheater of M=2.28 (SD =1.37). When testing all the ratings from both Studies 1 and 2 into a new repeated measures ANOVA, we did not obtain any significant main effect of Study nor an interaction with Musical Pieces (F<1), indicating that familiarity ratings did not differ across participants. However, a general tendency to be more familiar to Piazzolla than Dreamtheater was obtained (F(2,128)=2.98, p=.054).

#### 4.2.2 Regional fMRI activations from General Linear Model of BOLD response

##### Main effect of Aesthetic Value

As evidenced in Table 5 and Figure 3, the significant clusters (when corrected for multiple comparison with FWE) obtained for the main effect of Aesthetic Value were located bilaterally in the middle and superior temporal gyri and in the medial orbitofrontal cortex. Specifically, the mOFC was recruited by the contrast Beautiful>Ugly, an area involved in pleasure and reward processes. The same contrast also produced significant (p-value uncorrected) brain activations located in left middle frontal gyrus. The FWE-corrected bilateral (slightly right-predominant) temporal activations were instead to be ascribed only to the contrast Ugly>Beautiful (Figure 3).

**Figure 3.**
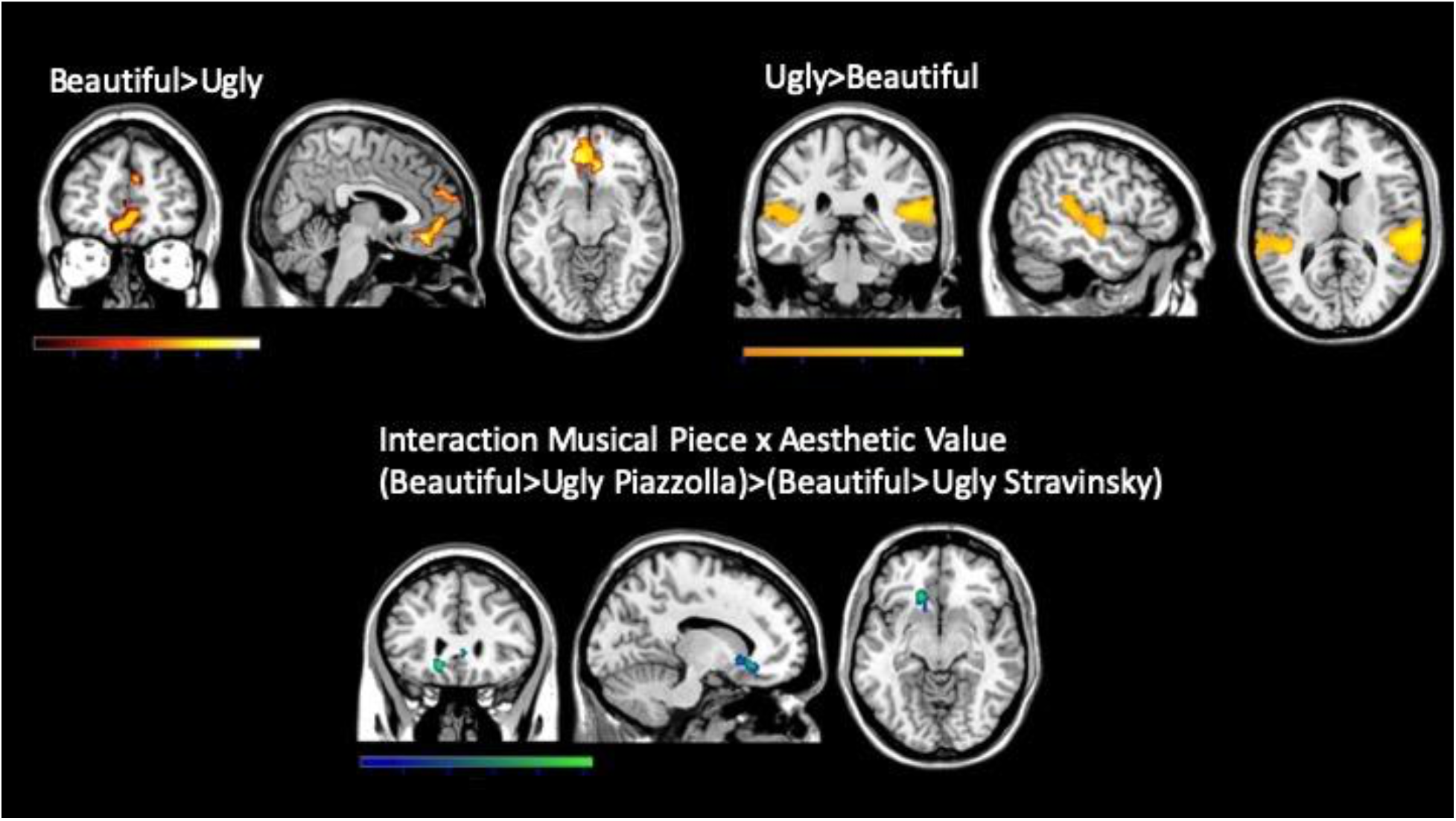
Top left: Brain activations in the medial orbitofrontal cortex related to the experience of the music passages evaluated most consistently by participants as beautiful. Voxels that survived to the statistical threshold of p < .001 uncorrected with an extent threshold of k=100 are shown. The colour scale illustrates the corresponding Z values. Top right: Brain activations in the superior and middle temporal gyri related to contrast Ugly>Beautiful. Voxels reported were significantly activated using a p-value <.05 corrected for multiple comparisons (FWE). The color scale illustrates the corresponding Z values. Bottom: Brain activations in the medial orbitofrontal cortex related to the interaction Musical Piece X Aesthetic value. Voxels reported were significantly activated using a p-value <.05 corrected for multiple comparisons (FWE). The color scale illustrates the corresponding Z values.

In the GLM analysis including acoustic components as regressors of no interest (see Table 6), while the orbitofrontal cluster was quite preserved (p-value uncorrected) for the contrast Beautiful>Ugly, the temporal clusters for the Ugly>Beautiful contrast were not significant anymore, even when lowering the alpha thresholds.

##### Interaction Music*Beauty

As illustrated in Figure 3, the FWE-corrected interactions between factors Musical Piece and Aesthetic Value resulted in significant activations in a focal orbitofrontal cluster. In particular, Beautiful>Ugly in Piazzolla compared to Beautiful>Ugly in Stravinsky led to greater neural activity in mOFC (also when including the acoustic components in the analyses, but uncorrected). This suggests that mOFC activity mainly occurred when listening to the beautiful passages of Piazzolla.

##### Main effect of Musical Piece

The main effect of Musical Piece revealed increased activity in three clusters (FWE-corrected) within the bilateral superior temporal gyrus (Figure 4). These activations were better explained by the contrast Piazzolla>Stravinsky, whether its reciprocal did not produce any significant activation. In the GLM analysis including acoustic components are regressors of no interest (Figure 4), the stronger activation in left STG remained significant (FWE-corrected) for the contrast Piazzolla > Stravinsky.

**Figure 4.**
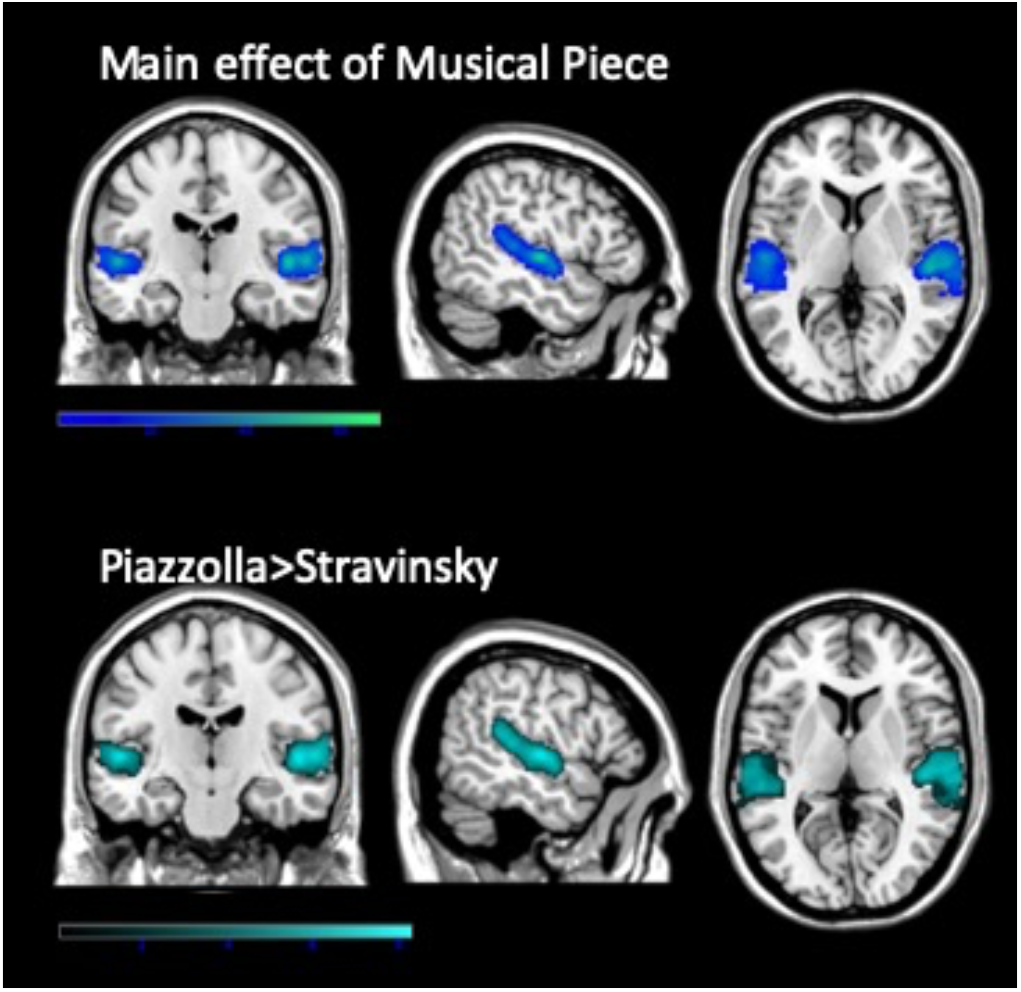
Top: Brain activaitons in the superior temporal gyri related to the main effect of Musical piece. Bottom: Adios Nonino>Rite of Spring. Increased brain activity in the superior temporal gyri during listening to Adios Nonino as opposed to Rite of Spring, with acoustic features regressed out from the analysis. Voxels reported in the illustration were significantly activated using a p-value <.001 uncorrected with and extent threshold of k=100. The colour scale illustrates the corresponding Z values.

#### 4.2.3 Dynamic Causal Modelling

##### Effective connectivity during beautiful musical passages

We analyzed the strength of extrinsic connections between bilateral STG and OFC and intrinsic (inhibitory) connections within each region. As visible from Figure 5, Bayesian model comparison of model evidences at the first level (Penny et al. 2010) revealed that the cortical network with changes in both extrinsic and intrinsic connections was the most frequent across subjects (Posterior model probability > 0.85 and protected exceedance probability > 0.98). Hence, the winning model was the first full one. This was confirmed by a Bayesian model comparison of the model evidences at the second level under parametric empirical Bayes (Posterior model probability > 0.99). Within this network, we found an increase in feedforward connectivity from left STG to OFC (Posterior probability 0.98) and from right STG to OFC (Posterior probability 0.95). We also found an increase in the inhibitory connectivity within both left STG (Posterior probability > 0.99) and right STG (Posterior probability > 0.99). In other words, when subjects perceived the stimuli as beautiful, there was a consistent increase in connection strengths from bilateral STG to OFC and a concomitant increase in cortical inhibition within bilateral STG.

##### Effective connectivity during ugly musical passages

As evidenced in Figure 5, when subjects listened to ‘ugly’ musical passages, DCM showed a decrease in feedforward connectivity from right STG to OFC (Posterior probability 0.95), namely the winning model was the third reduced feedforward one. DCM also evidenced a decrease in the inhibitory connectivity within both left STG (Posterior probability > 0.99) and right STG (Posterior probability > 0.99). In other words, when subjects listened to ugly musical passages, there was a decrease in connection strength from right STG to OFC and a concomitant disinhibition within bilateral STG.

### 4.3 Study 3

As illustrated in Figure 6, the music composition experts provided ratings that significantly discerned the beautiful from the ugly passages of the musical pieces as identified in Study 1. The passages evaluated as beautiful or ugly differed on most of the aesthetic dimensions (Table 7), with the ‘ugly’ fragments rated as more atonal, innovative, agitating, fast, rhythmic and with a higher complexity in harmony, rhythm and execution when compared to the ‘beautiful’ excerpts, which received opposite evaluations on the other poles of the same dimensions.

**Figure 6.**
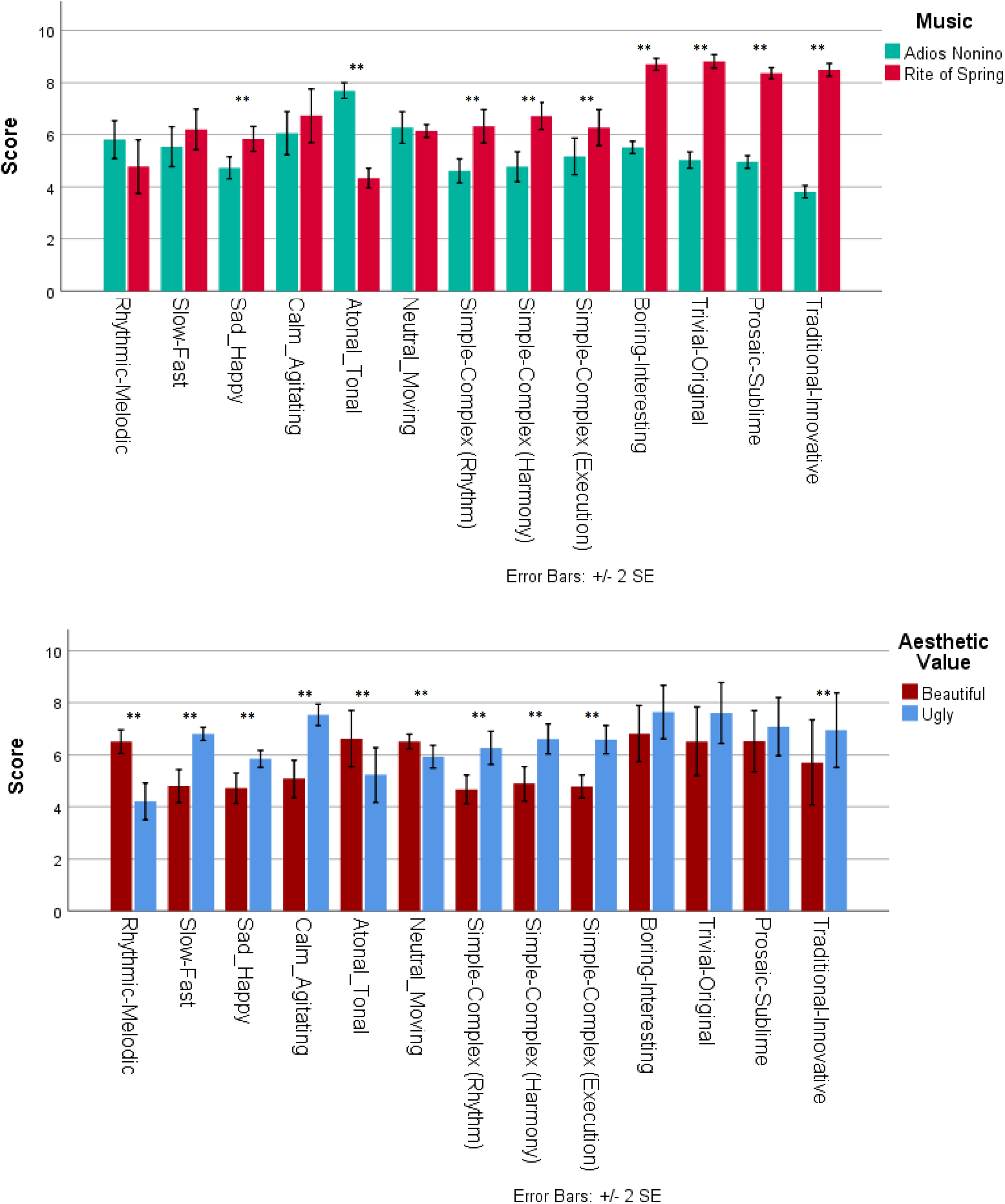
Questionnaire results. The two sets of histograms show the average scores and standard deviations of musical and affective features given by experts to each musical passage identified as beautiful or ugly from Study 1. The passages belonged to the pieces “Adios Nonino” by A. Piazzolla and “Rite of Spring” by I. Stravinsky. The musical (rhythmic-melodic, slow-fast, atonal-tonal, simple-complex rhythm, simple-complex execution, trivial-original, traditional-innovative) and affective (sad-happy, calm-agitating, neutral-moving, boring-interesting, prosaic-sublime) were given on a scale from 1 to 10. Lines and asterisks indicate the outcomes of the ANOVAs contrasting either the two pieces (top) or ugly vs. beautiful ratings (bottom).

Moreover, the ratings were idiosyncratic to each musical piece, as shown by the significant main effects of Musical Piece in the ANOVAs on the several aesthetic dimensions (Table 7), resulting from more tonal ratings for Piazzolla than Stravinsky, but more joyful, interesting, original, sublime, innovative and complex (in relation to harmony, rhythm and performative style) for Stravinsky over the simpler, sadder, and more boring, trivial, prosaic and traditional passages by Piazzolla as compared with Stravinsky. In Piazzolla, but not in Stravinsky, the beautiful passages were evaluated as more moving than the ugly ones.

## 5. DISCUSSION

In this research we aimed to identify the neural activity associated with the inter-subjective aesthetic appraisal of music during a realistic listening condition. Additionally, we aimed to understand the musical properties found to elicit highly agreed aesthetic responses. This was achieved by using an original three-step strategy. In a first study, we obtained continuous evaluations of beauty of three musical pieces by means of a motion sensor, for selecting the musical passages that were consistently rated with a positive or negative valence across participants. In a second study, neuroimaging data during naturalistic music listening were obtained in a separate session from a new group of adult participants; the most consistent intervals in the behavioral ratings from the first study informed the neuroimaging analysis. Findings revealed focal activity and modulation of effective connectivity in two brain structures by aesthetic value of music: the medial orbitofrontal cortex and the superior temporal gyrus. In a third study, the selection of the most consistent beautiful and ugly passages in the music were further rated by music composition experts on several musical and emotional dimensions for inquiring whether any specific feature can be consistently associated with beauty evaluations of music in the general Western population, irrespectively of subjective listening biography or personal characteristics. Questionnaire information showed that when the music passages shifted toward an evaluation of beauty they were more tonal, traditional, melodic, sad, calm, slow and simple which all together recall a melancholic and low arousing emotional state, similar to the emotions of ‘tenderness’ and ‘peacefulness’ in the GEMS model by (Zentner, Grandjean, and Scherer 2008).

The consistency of current findings is grounded in the protocol adopted for this naturalistic study, where three different studies were conducted with three separate samples of Western participants, having a broad educational and cultural background and ranging in age from 18 to 57 years. The results converged in identifying a common neural mechanism for subjective musical beauty relying on neural communication between auditory and orbitofrontal regions, as well as objective features related to musical structure and affective connotations that are consistently eliciting positive aesthetic evaluations in the general (Western) population. While being a unique neuroscientific exploration on positive and negative aesthetic responses to music, this first research calls for further investigations that would generalize the findings to non-Western listeners and account for the full variety of music from different genres and played with different instruments up to even songs and lyrics.

### 5.1 Musical beauty and the medial orbitofrontal cortex

Our study represents a contribution to the scarce neuroimaging literature on aesthetic responses to music. Most previous studies focused on specific emotions (e.g., Brattico et al. 2011; Bogert et al. 2016), sensory pleasure (Blood et al. 1999), and processing of musical features (e.g., tonality, timbre, rhythm: Proverbio, Orlandi, and Pisanu 2016; Alluri et al. 2012; Zatorre and Belin 2001; Platel et al. 1997). Few recent investigations showed the involvement of the reward circuit and its connectivity with auditory and inferofrontal regions during enjoyment of instrumental music. To our knowledge only one study, thus far, addressed the question of neural mechanisms for attributing beauty value to music (Ishizu and Zeki 2011). This contrasts with the wide interest for understanding the neural correlates of this common type of judgment in visual cognitive neuroscience (Pearce et al. 2016; Tiihonen et al. 2017).

The most striking finding of our study is the focal orbitofrontal activity in the medial OFC in response to musical beauty, when considered irrespectively of the subjective reactions and individual listening biographies. The same focal activity was found by (Ishizu and Zeki 2011), although in their experiment the paradigm was far from naturalistic and involved a behavioral task and interruptions of aesthetic contemplation. The OFC activity there was obtained both in response to beautifully rated musical passages as well as beautiful paintings, hence, irrespectively of the sensory modality that originated the beauty experience. Accordingly, they proposed a causal link between OFC activity and the perception of beauty, namely that the neuronal firing in this region would cause the attribution of beauty qualities to any given stimulus. Moreover, many other investigators have associated the broader region of the orbitofrontal cortex, including several citoarchitectonic areas (Brodmann areas 10, 11, 12, 32 and 25) to different types of pleasurable stimuli, proposing a hierarchical organization with primary pleasures, such as sex and food to lateral areas and secondary or abstract pleasures such as money and music to more medial areas (Brown et al. 2011; Kawabata and Zeki 2004; Vartanian and Goel 2004; Stark, Vuust, and Kringelbach 2018). In studies focusing only on musical pleasure, OFC activity often accompanies listening to pleasant music, whether its pleasantness is strictly driven by acoustic features, such as consonance or harmonicity and absence of roughness and beats in pleasant and happy sounds (Bogert et al. 2016; Koelsch 2014), and independently from the negative valence of musical appraisal, such as the enjoyment of songs including sad ones (Elvira Brattico et al. 2015; Salimpoor et al. 2011; A. J. Blood and Zatorre 2001). Indeed, the OFC role for experiencing musical emotions in general has been confirmed by a recent meta-analysis (Koelsch 2014). In sum, the here-obtained activity in the medial OFC is usually interpreted as reflecting the reward value of the presented artworks or aesthetic stimuli, probing the relationship with reward, pleasure and judgment.

In turn, we did not observe any significant (or even below-significant threshold) activation in other areas of the reward system that have been implied in other studies of musical aesthetic responses (specifically, pleasure) such as ventral striatum, cingulate cortex and ventromedial prefrontal cortex (for reviews, see Robert J. Zatorre and Salimpoor 2013; Reybrouck, Vuust, and Brattico 2018; García-Prieto, Pereda, and Maestú 2016). This might be related to the original goal of the study, namely to search for the invariant neural features associated to musical beauty, which are independent of sensory processing and related pleasurable responses to sounds and independent of the listener’s background. In other words, when beauty is studied as an abstract property distinct from sensory pleasure, as a value that is cognitively attributed to stimuli based on their formal, structural, invariant aspects, then the supramodal region of the orbitofrontal cortex seems to emerge as its central hub. Similar findings are obtained in studies of visual neuroaesthetics, where the orbitofrontal region is the main center of value attribution to an artwork (Pearce et al. 2016; Chatterjee and Vartanian 2016). Notably, the involvement of mOFC seems predominant especially during the perception of beautiful art or music stimuli, namely not when an explicit judgment is provided. For instance, in a study where mediofrontal activity was obtained in response to aesthetic enjoyment of familiar music by (Pereira et al. 2011), participants judged the preference and familiarity of musical excerpts in a behavioral session and then were asked to passively listen to them in the fMRI scanner. In turn, when a conscious decision is provided by means of a behavioral response like a button press, the cognitive dorsolateral prefrontal cortex (DLPFC) seems to be recruited. Indeed, several researchers (Ticini 2017; Marcos Nadal et al. 2008; Cela-Conde et al. 2013; Wallis and Miller 2003) converge in suggesting that the reward signal reaches quickly the orbital area and is forwarded to the dorsolateral prefrontal cortex for guiding the forthcoming behavioral response (a goal-directed action).

Our mOFC findings were mainly driven by Piazzolla stimulus, “Adios Nonino”, suggesting by reverse inference that the piece was more effective than the one by Stravinsky to be associated with beauty judgments. This is resonant with previous findings obtained by our lab with the naturalistic free-listening paradigm (e.g., Liu et al. 2017). Of all the stimuli used, “Adios Nonino” produced the most statistically powerful brain signal especially when linear regression models were considered. Such observation might be a consequence of the acoustic variability in the musical piece compared with others, allowing less neuronal adaptation (Haumann et al. in press) and the availability of better times series regressors when acoustic features have been used for the analysis. Indeed, when listening to Piazzolla, participants showed also a stronger neurometabolic signal in the left superior temporal gyrus, indicating that the music piece enhanced activity in the auditory areas yet excluding the role of acoustic features. This phenomenon has been proved to happen when listening to consonant, liked music (e.g., Koelsch, Skouras, and Lohmann 2018; Brattico et al. 2015). Aesthetic appraisals engage those regions of the brain that underlie sensation and perception relative to the modality through which the object is presented (Nadal et al. 2008). A simpler explanation of the auditory-cortex findings for Adios Nonino might be the difference in the statistical power between the conditions, although the difference between them amounted to few volumes (Piazzolla: 98 sec beautiful, 95 ugly; Stravinski: 74 sec beautiful, 82 sec ugly). On the other hand, while the two stimuli were similarly liked by participants, the preference for the Stravinsky piece correlated with musical expertise. Hence, it is possible that the heterogeneity of the participants’ musical background might have played a role in the finding. The experts’ preference for Stravinsky is understandable considering their analytical knowledge of the work of art and its aesthetic value. Future studies should systematically test whether the expertise for a musical genre modulates OFC activity to beautiful music.

The questionnaire data of Study 3 collected from 12 music-composition experts further elucidates how formal, structural aspects of music can invariantly drive aesthetic judgments in a Western general population. According to the experts, the passages of music that were most consistently evaluated in Study 1 as beautiful, for both musical pieces, were more melodic, calm, sad, slow, tonal, traditional and simple than the ones that were consistently judged as ugly. The music piece “Adios Nonino” was rated by the experts as more tonal, sad, boring, prosaic, trivial and simpler than the other one. Moreover, its beautiful music passages were evaluated more moving than the others. These features all together in our opinion recall a melancholic, peaceful and low arousing emotional state, similar to the aesthetic emotions of ‘tenderness’ and ‘peacefulness’ in the GEMS model by (Zentner, Grandjean, and Scherer 2008). The findings seemed to confirm that the most statistically common features in music across the world consist of discrete pitch organisation and melodic patterns, leading to a general preference for consonant, tonal music (e.g., Mencke et al. 2018; Butler and Daston 1968; Savage et al. 2015). Hence, our results suggest that, beyond the variability in subjective aesthetic judgements, a high degree of consensus can be found in the general population on those passages in the music that evoke emotions of tenderness and nostalgia. These passages are those that are intersubjectively interpreted as more “beautiful” and that are associated with a higher activation in mOFC, as indeed observed also in other neuroimaging studies focusing on musical emotions rather than beauty (Trost et al. 2012; Khalfa et al. 2005; Barrett and Janata 2016). Our findings bring a first brick of evidence on the neural mechanisms involved during aesthetic appreciation and, more specifically, during the experience of musical beauty. Future studies should go at least in two directions. On the one hand, they should determine if the same consensus on the aesthetic value of the musical passages individuated by us can be found among non-Western listeners. Moreover, future studies should determine how much is the amount of variability in the aesthetic judgement that can be explained by individual differences in listening biography and musical expertise. Indeed, a recent study showed how individual profiles that can be mapped into distinct clusters modulate the preference for specific acoustic features (Güçlütürk, Jacobs, and van Lier 2016).

### 5.2 Negative appraisal of music and the auditory cortex

In the present study, the musical passages that were consistently rated as ugly were associated with bilateral brain activity in the superior temporal gyrus (STG), although with a predominance of the right hemisphere. This brain activity was significantly reduced when the sensory processing of acoustic features were pruned from the neurometabolic signal by including as regressors of no interest in the general linear model the computationally extracted timbral, tonal, and rhythmic parameters (Fullness, Brightness, Timbral Complexity, Key Clarity, Pulse Clarity, and Activity; see Alluri et al. 2012). Hence, it seems that the ugly passages were characterized by a higher acoustic complexity as well as by a larger neuronal recruitment of brain regions dedicated to auditory processing, particularly the supratemporal cortex. Our results hence indicate that superior temporal gyrus activity during listening to ugly musical passages was related to the acoustic and aesthetic features characterizing them and requiring more demanding sensory processing. Indeed, (Berlyne 1971) and more recently exponents of the probabilistic account of music cognition (Huron 2006; Gebauer, Kringelbach, and Vuust 2012; Brattico 2019) proposed that an inverted U-shaped curve reflects a general relationship between aesthetic appreciation and structural complexity. The passages of music consistently evaluated as ugly were described by composers in Study 3 as more complex than the beautiful ones in regard of harmonic, executive and rhythmic complexity; more specifically, they were described as agitating, joyful, fast, atonal, innovative, complex and rhythmic.

Literature on the counterpart of aesthetic experience of music, i.e., the experience of disliked and negatively valenced musical stimuli, is not very abundant and does not yet present such a clear picture on the neural implications. For instance, Ishizu and Zeki (2011) in their study focusing on cross-modal (visual and musical) beauty did not find any area that was specifically involved with listening to ugly music excerpts. Typically, the negative experiences of music have been studied by using dissonant or rough music, that has been found to be perceived as more unpleasant than consonant music among people of different music cultures (Fritz et al., 2009) and age (Zentner & Kagan, 1998). In neuroimaging studies, dissonant music has been associated with activation in several limbic regions including the amygdala, hippocampus and parahippocampal gyrus (right), as well as in the precuneus and temporal poles (e.g., Blood et al. 1999; Koelsch et al. 2006). Moreover, the left auditory cortex seems to be preferentially recruited during listening to dissonant music (Proverbio, Orlandi, and Pisanu 2016). These features resemble the aesthetic emotions of joy and tension of the GEMS model by Zentner, Grandjean, and Scherer (2008) and are characteristic of a high-arousal and excited emotional state. Agitation and high arousing emotions (tension, excitement, anxiety) have been associated in the literature with activation of the amygdala and sensory and motor areas (Koelsch 2014). For instance, Trost et al. (2012) showed that high arousal emotions, both positively and negatively valenced such as tension, power and joy, also correlated with stronger brain activity in the superior temporal gyrus, among other areas. Proverbio et al. (2015) further proved that listening to atonal music reduced heart rate (fear bradycardia) and increased blood pressure, suggesting that it induces a parasympathetic response of increased arousal and alertness. In line with this, our findings indicate that the reaction to ugly music might be influenced by its auditory complexity and by its intrinsic threatening and agitating properties. Yet, as previously mentioned, future studies should disclose the role of musical expertise in modulating the level of dissonance that can be processed and aesthetically appreciated by the human cognitive system.

### 5.3 Effective connectivity between auditory and orbitofrontal cortices

In the past decade, the brain physiology is increasingly understood as a complex network system of nodes interacting dynamically with each other. Studies of brain connectivity and networks are exponentially accumulating covering all the aspects of human cognitive functions, except perhaps aesthetic experience. After an initial proposal (Lang et al. 1993), the idea that brain networks might be modulated during an aesthetic experience was put forward by several scientists (Cela-Conde et al. 2004; Jacobsen 2014; Ramachandran and Hirstein 1999; Brattico, Bogert, and Jacobsen 2013; Pelowski et al. 2017), all pointing out the dynamic interaction of perceptual, attentional and emotional systems during an aesthetic experience. (Gallese and Di Dio 2012) further proposed that during an aesthetic experience the beholder “perceives–feels–senses” an artwork, which activates sensorimotor, emotional, and cognitive mechanisms. Even Zeki (2013) proposed that the flow from sensory cortices to orbital areas would dictate a beauty verdict. In spite of these theoretical proposals, it is not until very recently that empirical analyses of brain connectivity in relation to artworks appeared (e.g., Belfi et al. 2019; Wilkins et al. 2014).

In music, a recent review (Reybrouck, Vuust, and Brattico 2018) listed 12 studies of functional connectivity during listening to pleasurable music, and of them 8 included behavioral ratings of positive aesthetic responses to music. The findings converge in identifying a reward brain network including ventral striatum (specifically, the nucleus accumbens) and medial OFC in connection with auditory frontotemporal areas as the circuit responsible for attributing motivational value to music, quantified as the intention to pay smaller or larger amounts of dollars for purchasing a tune (Salimpoor et al. 2013). Even the lack of hedonic responses to music seems to depend on weaker connections between supratemporal regions and ventral striatum (Martínez-Molina et al. 2016). Very recently, Fasano and colleagues (2020) further noticed higher probability of connectivity patterns within orbital regions in the brains of pre-adolescent children while listening to pleasurable music as opposed to a rest condition. These sparse studies on functional connectivity during musical appreciation though, and during visual aesthetic contemplation alike, have not yet determined the temporal interaction between brain structures, thus missing to capture the causal dependence of neural communication between interconnected brain structures.

Only a previous study by Kumar et al. (2012) applied DCM to understand how affective reactions to sounds are encoded in the brain. Findings showed that a cortico-amygdalar pathway is needed for attributing negative valence to aversive sounds (those sounds that are characterized by a frequency range between 2500 and 5500 Hz, and temporal modulations in the range 1–16 Hz). In particular, while acoustic features of aversive sounds modulated the feedforward connections from auditory cortex to amygdala, valence attribution modulated the connections from amygdala to auditory cortex. For revealing the temporal dependency of connectivity patterns representing the experience of beauty during a naturalistic listening of music, we produced four dynamic causal models with DCM. Each aimed to predict the attribution of beauty versus ugliness to musical passages based on different interaction patterns between the STG and OFC clusters found active in the GLM analysis: 1) both extrinsic (excitatory) and intrinsic (inhibitory) connections between auditory and orbital areas; 2) only extrinsic (excitatory) connections between auditory and orbital areas; 3) only feedforward (excitatory) connections; 4) no connections. The first full model was most probable during passages rated as more beautiful whereas the third reduced, feedforward model predicted best the neural communication during listening to those passages rated as less beautiful (ugly). Hence, during perception of beautiful musical passages, the connections from bilateral auditory cortices to orbital areas increased concurrently with a bilateral cortical inhibition of bilateral supratemporal regions. In turn, the opposite happened during listening to ugly musical passages, since connections from the right supratemporal region to orbitofrontal cortex decreased while bilateral auditory cortices were active. Overall, our analysis evidenced how the neural interplay between orbital area and auditory cortex either ensues into a positive aesthetic experience or into a negative one: a positive aesthetic appraisal occurs when the auditory system inhibits itself and send forward neural signal to orbitofrontal areas whereas in the case of the observed negative appraisal the neural signal remains within bilateral auditory cortices and coupling from right supratemporal region to orbital area is inhibited.

Our findings resonate with rare observations of neural chronometry for affective sounds, as obtained by applying neural source modelling to magnetoencephalography (MEG), a neuroimaging modality allowing very fine temporal resolution. In other words, directionality of brain activations can be also determined in terms of phase and time delays between neural firings. For instance, aversive reactions to evolutionarily salient sounds such as infant vocalisations were found to occur as early as 50ms from sound onset in the periaqueductal gray (PAG) in the midbrain and to travel directly to OFC for valence attribution (Young et al. 2017). Within the realm of art-related responses, neural chronometry proposals of aesthetic processing have been outlined by Brattico, Bogert, and Jacobsen (2013) and Pelowski et al. (2017) on the grounds of neurophysiological and neuroimaging findings. These proposals, in line with our DCM findings, suggest that positive aesthetic judgments are represented in OFC and DLPFC areas, and occur only after initial processing stages, including impression formation in sensory cortices and early and later emotional reactions in the limbic system.

## 6. CONCLUSIONS

The present work aimed at determining the neural mechanisms associated with the subjective appraisal experience of musical beauty in the general population, and its relationship with the objective musical features, by using a convergence of neural, behavioral and questionnaire measures. The neural mechanisms involved in the subjective experience of musical beauty among a sample of Western listeners with a heterogeneous listening history and context were found in two brain structures: the mOFC and the bilateral STG. During the experience of musical beauty, the regional STG activity became inhibited and coupled with mOFC, whereas during perception of musical ugliness, regional STG activity became stronger and decoupled from mOFC. The identified neural mechanisms as obtained in the neuroimaging study were paralleled by a set of musical and emotional features that discerned the beautiful from the ugly passages on the basis of music-composition experts’ ratings. Hence, in line with previous findings from both music and other art domains, we can conclude that the mOFC represents a brain hub for the inter-subjective experience of beauty in music, such that it occurs with the consensual identification of emotional features of sadness, tenderness and pathos, and of musical features of simplicity, tonality and slower tempo in music passages by people of varying degrees of musical expertise. In turn, the aesthetic attribution of low beauty or ugliness in music is associated – across participants with different musical expertise, musical pieces and measurement modalities –with more complex processing, and more arousing and agitating musical content.

## 7. ACKNOWLEDGMENTS

The study could not be possible without the help of research assistants and collaborators both at the stage of data collection and analysis: Brigitte Bogert, Houra Saghafifar, Dr. Marina Kliuchko, Benjamin Gold, Taru Numminen-Kontti, Mikko Heimölä, Dr. Jyrki Mäkelä, Toni Auranen, Marita Kattelus, Dr. Mari Tervaniemi, Dr. Teppo Särkämö, Dr. Vinoo Alluri, Leonardo Bonetti. The Center for Music in the Brain (MIB) is funded by the Danish National Research Foundation (DNRF project 117).

## SUPPLEMENTARY MATERIAL

### Stimuli

The version of “Adios Nonino” used for the study was picked from the CD “The Lausanne Concert” (BMG Music, 1993). The piece was recorded live at MAD (Moulin a Danses), Lausanne, Switzerland on November 4^th^ 1989, as part of the Sexteto European tour (1989-1990). The duration of the track was 08’07’’968 and it was performed by Astor Piazzolla (bandoneon), Horacio Malvicino (guitar), Carlos Nozzi (violoncello), Angel Ridolfi (double bass, upright bass), Daniel Binelli (bandoneon) and Gerardo Gandini (piano).

The version of “The Rite of Spring”was extracted from the CD “Stravinsky: The Rite of Spring/Scriabin: The Poem of Ecstasy” (Philips, 2001). The piece was performed by the Orchestra of the Kirov Opera of St. Petersburg and directed by Valerij Gergiev. The excerpts of the three episodes (Introduction: 00:05-03:23, The Augurs of Spring: 0-03:12, Ritual of Abduction: 0-01:16) were comprised together for a total duration of 07’’ 47’ 243.

